# Rapid encoding of task regularities in the human hippocampus guides sensorimotor timing

**DOI:** 10.1101/2021.08.03.454928

**Authors:** Ignacio Polti, Matthias Nau, Raphael Kaplan, Virginie van Wassenhove, Christian F. Doeller

**Affiliations:** Kavli Institute for Systems Neuroscience, Centre for Neural Computation, The Egil and Pauline Braathen and Fred Kavli Centre for Cortical Microcircuits, Jebsen Centre for Alzheimer’s Disease, Norwegian University of Science and Technology, Trondheim, Norway; Max-Planck-Institute for Human Cognitive and Brain Sciences, Leipzig, Germany; Department of Basic Psychology, Clinical Psychology, and Psychobiology, Universitat Jaume I, Castellón de la Plana, Spain; CEA DRF/Joliot, NeuroSpin; INSERM, Cognitive Neuroimaging Unit; CNRS, Université Paris-Saclay, Gif-Sur-Yvette, France; Institute of Psychology, Leipzig University, Leipzig, Germany

## Abstract

The brain encodes the statistical regularities of the environment in a task-specific yet flexible and generalizable format. Here, we seek to understand this process by converging two parallel lines of research, one centered on sensorimotor timing, and the other on cognitive mapping in the hippocampal system. By combining functional magnetic resonance imaging (fMRI) with a fast-paced time-to-contact (TTC) estimation task, we found that the hippocampus signaled behavioral feedback received in each trial as well as performance improvements across trials along with reward-processing regions. Critically, it signaled performance improvements independent from the tested intervals, and its activity accounted for the trial-wise regression-to-the-mean biases in TTC estimation. This suggests that the hippocampus supports the rapid encoding of temporal context even on short time scales in a behavior-dependent manner. Our results emphasize the central role of the hippocampus in statistical learning and position it at the core of a brain-wide network updating sensorimotor representations in real time for flexible behavior.

## Introduction

When someone throws us a ball, we can anticipate its future trajectory, its speed and the time it will reach us. These expectations then inform the motor system to plan an appropriate action to catch it. Generating expectations and planning behavior accordingly builds on our ability to learn from past experiences and to encode the statistical regularities of the tasks we perform. At the core of this ability lies a continuous perception-action loop, initially proposed for sensorimotor systems (e.g. Wolpert et al. (2011)), which is now at the heart of many leading theories of brain function including active inference (Friston et al., 2016), predictive coding (Huang & Rao, 2011) and reinforcement learning (Daw & Dayan, 2014).

Critically, the brain needs to balance three primary objectives to effectively guide behavior in a dynamic environment. First, it needs to capture the specific aspects of the task that inform the relevant behavior (e.g. the remaining time to catch the ball). Second, it needs to generalize from a limited set of examples to novel and noisy situations. This can be achieved by regularizing the currently encoded information based on past experiences (e.g. by inferring how fast previous balls flew on average). Third, the sensorimotor representations that guide the behavior need to be updated flexibly whenever feedback about our actions becomes available (e.g. when we catch or miss the ball), or when task demands change (e.g. when someone throws us a frisbee instead). Herein, we refer to these objectives as specificity, regularization and flexibility. While these are all fundamental principles underlying human cognition broadly, how the brain forms and continuously updates sensorimotor representations that balance these three objectives remains unclear.

Here, we approach this question with a new perspective by converging two parallel lines of research centered on sensorimotor timing and hippocampal-dependent cognitive mapping. Specifically, we test how the human hippocampus, an area implicated in memory formation on long time scales (days to weeks), may support the formation and flexible updating of sensorimotor representations even on short time scales (milliseconds to seconds). We do so by characterizing in detail the relationship between hippocampal activity and behavioral performance in a fast-paced timing task, which is traditionally believed to be hippocampal-independent. We propose that the capacity of the hippocampus to encode statistical regularities of our environment (Behrens et al., 2018; Momennejad, 2020; Whittington et al., 2020) situates it at the core of a brain-wide network balancing specificity vs. regularization in real time as the relevant behavior is performed.

An optimal behavioral domain to study these processes is sensorimotor timing (Gershman et al., 2014; Petter et al., 2018). This is because prior work suggested that timing estimates indeed rely on the statistics of prior experiences (Wolpert et al., 2011; Jazayeri & Shadlen, 2010; Acerbi et al., 2012; Chang & Jazayeri, 2018). Crucially, however, timing estimates are not always accurate. Instead, they directly reflect the trade-off between specificity and regularization, which is expressed in systematic behavioral biases. Estimated intervals regress towards the mean of the distribution of tested intervals (Jazayeri & Shadlen, 2010), a well-known effect that we will refer to as the regression effect (Petzschner et al., 2015). The regression effect suggests that the brain encodes a probability distribution of possible intervals rather than the exact information obtained in each trial (Wolpert et al., 2011). Timing estimates therefore depend not only on the interval tested in a trial, but also on the temporal context in which they were encountered (i.e., the intervals tested in all othertrials). This likely helps to predict future scenarios, to adapt behavior flexibly and to generalize to novel or noisy situations (Jazayeri & Shadlen, 2010; Acerbi et al., 2012; Roach et al., 2017).

Importantly, the hippocampus proper codes for time and temporal context on various scales (Howard, 2017) and it has been shown to process behavioral feedback in decision-making tasks (Shohamy & Wagner, 2008), pointing to a role in feedback learning. Moreover, the hippocampal formation has been implicated in encoding the latent structure of a task along with the individual features that were tested (Kumaran, 2012; Schlichting & Preston, 2015; Schapiro et al., 2017; Wikenheiser et al., 2017; Behrens et al., 2018; Schuck & Niv, 2019; Whittington et al., 2020; Peer et al., 2021), providing a unified account for its many proposed roles in navigation (Burgess et al., 2002), memory (Schiller et al., 2015; Eichenbaum, 2017) and decision making (Kaplan et al., 2017; Vikbladh et al., 2019). We propose that a central function of the human hippocampus is to encode the temporal context of stimuli and behaviors rapidly, and that this process manifests as the behavioral regression effect observed in time estimation and other domains (Petzschner et al., 2015). This puts the hippocampus at the core of a brain-wide network solving the trade-off between specificity and regularization for flexible behavior by continuously updating sensorimotor representations in a feedback-dependent manner. Using functional magnetic resonance imaging (fMRI) and a senso-rimotor timing task, we here test this proposal empirically.

## Results

In the following, we present our experiment and results in four steps. First, we introduce our task, which built on the estimation of the time-to-contact (TTC) between a moving fixation target and a visual boundary, as well as the behavioral and fMRI measurements we acquired. On a behavioral level, we show that participants’ timing estimates systematically regress towards the mean of the tested intervals. Second, we demonstrate that hippocampal fMRI activity and functional connectivity tracks the behavioral feedback participants received in each trial, revealing a link between hippocampal processing and timing-task performance. Third, we show that this hippocampal feedback modulation reflects improvements in behavioral performance over trials. We interpret this activity to signal the updating of task-relevant sensorimotor representations in real time. Fourth, we show that these hippocampal updating signals were independent of the specific interval that was tested and reflected the magnitude of the behavioral regression effect in each trial.

Notably, for each of the hippocampal main analyses, we also performed whole-brain voxel-wise analyses to uncover the larger brain network at play. We found that in addition to the hippocampus, regions typically associated with timing and reward processing signaled sensorimotor updating in our task, particularly the striatum. Follow-up analyses further revealed a striking distinction in TTC-specific and TTC-independent updating signals between striatal sub-regions. We conclude by discussing the potential neural underpinnings of these results and how the hippocampus may contribute to solving the trade-off between task specificity and regularization in concert with this larger brain network.

### Time-to-contact (TTC) estimation task

We monitored whole-brain activity using fMRI with concurrent eye tracking in 34 participants performing a TTC task. This task offered a rich behavioral read-out and required sustained attention in every single trial. During scanning, participants visually tracked a fixation target, which moved on linear trajectories within a circular boundary. The target moved at one of four possible speed levels and in one of 24 possible directions (Fig. 1A, similar to Nau et al. (2018a)). The sequence of tested speeds was counterbalanced across trials. Whenever the target stopped moving, participants estimated when the target would have hit the boundary if it had continued moving. They did so while maintaining fixation, and they indicated the estimated TTC by pressing a button. Feedback about their performance was provided foveally and instantly with a colored cue. The received feedback depended on the timing error, i.e. the difference between objectively true and estimated TTC (Figs. 1B), and it comprised 3 levels reflecting high, middle and low accuracy (Fig. 1C). Because timing judgements typically follow the Weber-Fechner law (Rakitin et al., 1998), the feedback levels were scaled relative to the ground-truth TTC of each trial. This ensured that participants were exposed to approximately the same distribution of feedback at all intervals tested (Figs. 1C, S1B). After a jittered inter-trial interval (ITI), the next trial began and the target moved into another direction at a given speed. The tested speeds of the fixation target were counterbalanced across trials to ensure a balanced sampling within each scanning run. Because the target always stopped moving at the same distance to the boundary, matching the boundary’s retinal eccentricity across trials, the different speeds led to four different TTCs: 0.55, 0.65, 0.86 and 1.2 seconds. Each participant performed a total of 768 trials. Please see Methods for more details.

**Figure 1:**
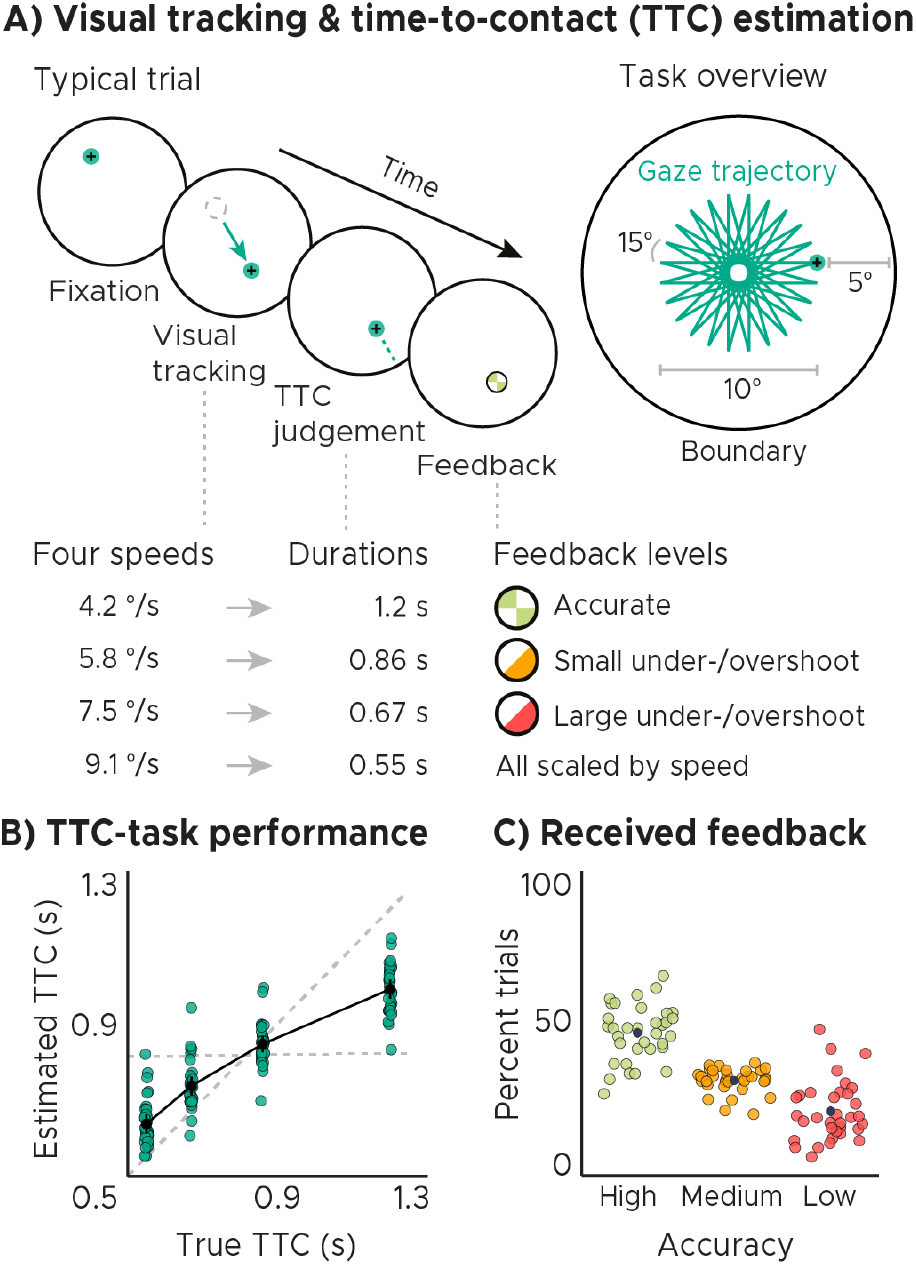
Visual tracking and Time-To-Contact (TTC) estimation task. A) Task design. In each trial during fMRI scanning, participants fixated a target (phase 1), which started moving at one of 4 possible speeds and in one of 24 possible directions for 10° visual angle (phase 2). After the target stopped moving, participants kept fixating and estimated when the fixation target would have hit a boundary 5° visual angle apart (phase 3). After pressing a button at the estimated TTC, participants received feedback (phase 4) according to their performance. Feedback was scaled relative to target TTC. B) Task performance. True and estimated TTC were correlated, showing that participants performed the task well. However, they overestimated short TTCs and underestimated long TTCs. Their estimates regressed towards the grand-mean of the TTC distribution (horizontal dashed line), away from the line of equality (diagonal dashed line). C) Feedback. On average, participants received high-accuracy feedback on half of the trials (also see Fig. S1B). BC) We plot the mean and SEM (black dots and lines) as well as single-participant data as dots. Feedback levels are color coded.

Analyzing the behavioral responses revealed that participants performed the task well and that the estimated and true TTCs were tightly correlated (Fig. 1B; Spearman’s *rho* = 0.91, *p* = 2.2*x*10^-16^). However, participants’ responses were also systematically biased towards the grand mean of the TTC distribution (0.82 seconds), indicating that shorter durations tended to be overestimated and longer durations tended to be underestimated. We confirmed this in all participants by examining the slopes of linear regression lines fit to the behavioral responses (Fig. S1C). These slopes differed from 1 (veridical performance; Fig. 1 B, diagonal dashed line; one-tailed one-sample *t* test, *t*(33) = −19.26, *p* = 2.2*x*10^-16^, *d* = −3.30, *CI* : [−4.22, −2.47]) as well as from 0 (grand mean; Fig. 1 B, horizontal dashed line; one-tailed one-sample *t* test, *t*(33) = 21.62, *p* = 2.2*x*10^-16^, *d* = 3.71, *CI* : [2.79,4.72]) and clustered at 0.5. Moreover, the slopes also correlated positively with behavioral accuracy across participants (Fig. S1D; Spearman’s *rho* = 0.794, *p* = 2.1*x*10^-08^), consistent with previous reports (Cicchini et al., 2012). Notably, the regression effect we observed in behavior has been argued to show that timing estimates indeed rely on the latent task regularities that our brain has encoded (e.g. Jazayeri & Shadlen (2010); Roach et al. (2017)). It may therefore reflect a key behavioral adaptation helping to regularize encoded intervals to optimally support both current task performance and generalization to future scenarios. In support of this, participants’ regression slopes converged over time towards the value of 0.5, i.e. the slope value between veridical performance and the grand mean (Fig. S1E; linear mixed-effects model with task segment as a predictor and participants as the error term, *F*(1) = 8.172, *p* = 0.005, *ϵ*^2^ = 0.08, *CI* : [0.01,0.18]). Visualizing the timing error over task segments and trials further showed that participants’ task performance improved over time (Fig. S1F; linear mixed-effects model with task segment as a predictor and participants as the error term, *F*(1) = 15.127, *p* = 1.3*x*10^-4^, *ϵ*^2^ = 0.06, *CI* : [0.02,0.11]), which suggests they were learning over the course of the experiment.

### Behavioral feedback predicts hippocampal activity

Importantly, sensorimotor updating is expected to occur right after the value of the performed action became apparent, which is when participants received feedback. As a proxy, we therefore analyzed how activity in each voxel reflected the feedback participants received in the previous trial. Using a mass-univariate general linear model (GLM), we modeled the three feedback levels with one regressor each (high, medium, low) plus additional nuisance regressors (see methods for details). We then contrasted the beta weights estimated for high-accuracy vs. low-accuracy feedback and examined the effects on group-level averaged across runs.

In both our regions-of-interest (ROI) analysis and a voxel-wise analysis, we found that hippocampal activity could indeed be predicted by the feedback participants received in the past trial (Figs. 2A,B). Higher-accuracy feedback resulted in overall stronger activity in the anterior section of the hippocampus (Figs. 2B, S2A; two-tailed one-sample *t* tests: anterior HPC, *t*(33) = −3.80, *p* = 5.9*x*10^-4^, *p_fwe_* = 0.001, *d* = −0.65, *CI* : [−1.03,-0.28]; posterior HPC, *t*(33) = −1.60, *p* = 0.119, *p_fwe_* = 0.237, *d* = −0.27, *CI* : [−0.62,0.07]). Moreover, the voxel-wise analysis revealed similar feedback-related activity in the thalamus and the striatum (Fig. 2A), and in the hippocampus when the feedback of the current trial was modeled (Fig. S3). Note that there was no systematic relationship between subsequent trials on a behavioral level (Fig. S1A; two-tailed one-sample *t* test; *t*(33) = 1.03, *p* = 0.312, *d* = 0.18, *CI* : [−0.17,0.52]; see methods for details) and that the direction of the effects differed across regions (Fig 2A), speaking against potential feedback-dependent biases in attention. Instead, these results are consistent with the notion that hippocampal activity signals the updating of task-relevant sensorimotor representations in real time.

**Figure 2:**
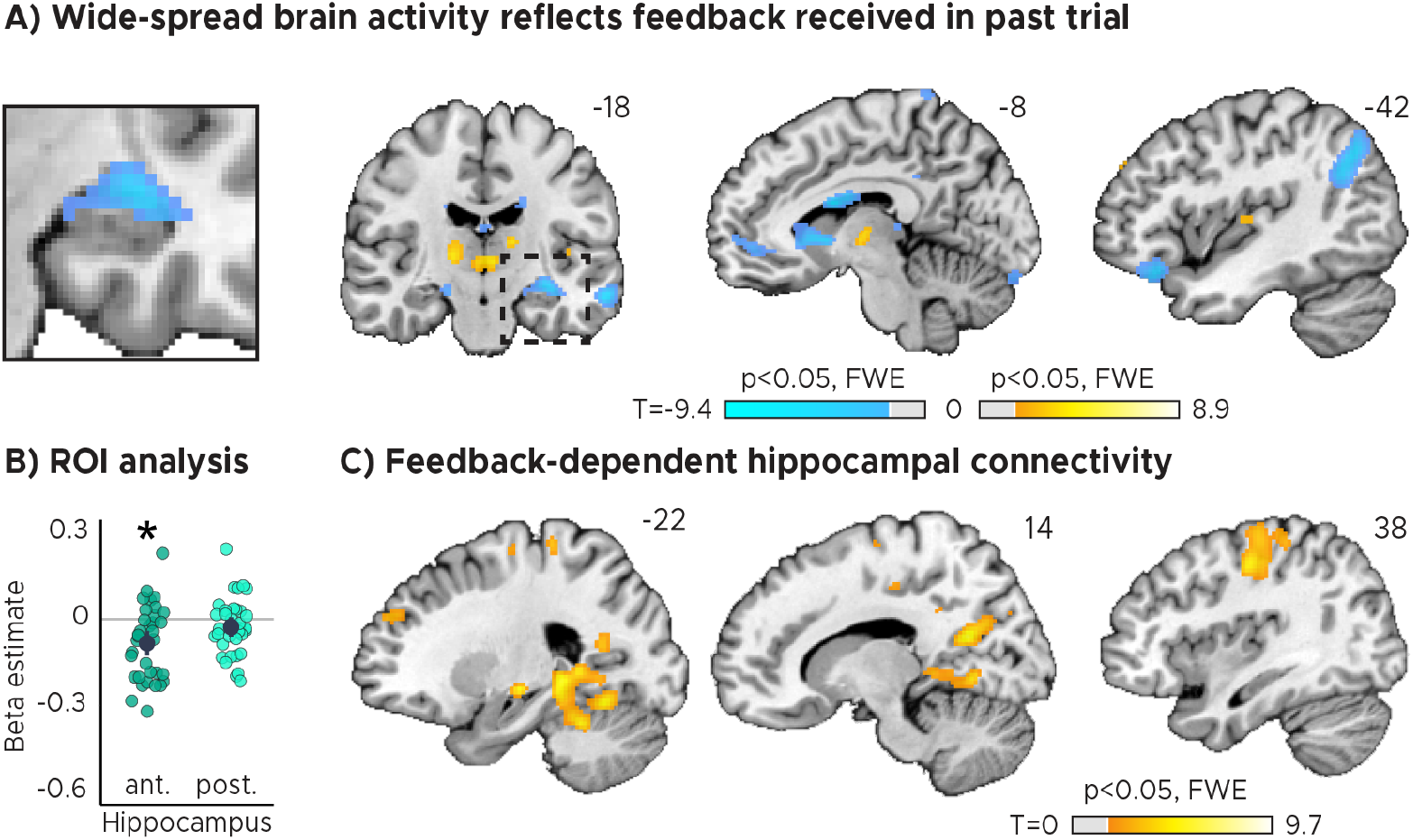
Feedback on the previous trial (n-1) modulates network-wide activity and hippocampal connectivity in subsequent trials (n). A) Voxel-wise analysis. Activity in each trial was modeled with a separate regressor as a function of feedback received in the previous trial. Insert zooming in on hippocampus added. B) Independent regions-of-interest analysis for the anterior (ant.) and posterior (post.) hippocampus. We plot the beta estimates obtained for the parametric modulator modeling trial-wise activity as a function of feedback in the previous trial. Negative values indicate that smaller errors, and higher-accuracy feedback, led to stronger activity. Depicted are the mean and SEM across participants (black dot and line) overlaid on single participant data (coloured dots). Activity in the anterior hippocampus is modulated by feedback received in previous trial. Statistics reflect p<0.05 at Bonferroni-corrected levels (*) obtained using a group-level two-tailed one-sample t-test against zero. C) Feedback-dependent hippocampal connectivity. We plot results of a psychophysiological interactions (PPI) analysis conducted using the hippocampal peak effects in (A) as a seed. AC) We plot thresholded t-test results at 1mm resolution overlaid on a structural template brain. MNI coordinates added. Hippocampal activity and connectivity is modulated by feedback received in the previous trial.

### Feedback-dependent hippocampal functional connectivity

Having established that hippocampal activity reflected feedback in the TTC task, we reasoned that its activity may also show systematic co-fluctuations with other task-relevant brain regions as well. To test this, we estimated the functional connectivity of a 4 mm radius sphere centered on the hippocampal peak main effect (x=-32, y=-14, z=-14) using a seed-based psychophysiological interaction (PPI) analysis (see methods). We reasoned that largertiming errors and therefore low-accuracy feedback would result in stronger updating compared to smaller timing errors and high-accuracy feedback, a relationship that should also be reflected in the functional connectivity between the hippocampus and other regions. We specifically tested this using the PPI analysis by contrasting trials in which participants performed poorly compared to those trials in which they performed well.

We found that hippocampal activity co-fluctuated with activity in the primary motor cortex, the parahippocampal gyrus and medial parietal lobe as well as the cerebellum (Fig. 2C). These cofluctuations were stronger when participants performed poorly in the previous trial and therefore when they received low-accuracy feedback. Combined with the previous analysis, this means that the absolute hippocampal activity scaled positively (Fig. 2A, B) and functional connectivity scaled negatively (Fig. 2C) with feedback valence.

### Hippocampal activity explains accuracy and biases in task performance

Two critical open questions remained. First, did the observed feedback modulation actually reflect improvements in behavioral performance over trials? Second, was the information that was learned specific to the interval that was tested in a given trial, likely serving task specificity, or was independent of the tested interval, potentially serving regularization? To answer these questions in one analysis, we used a GLM modeling activity not as a function of feedback received in the previous trial (Fig. 2), but as a function of the difference in feedback between trials (Fig. 3). Specifically, we modeled with two separate parametric regressors the improvements in TTC task performance across subsequent trials (regressor 1: TTC-independent updating) as well as the improvements over subsequent trials in which the same TTC interval was tested (regressor 2: TTC-specific updating). We again accounted for nuisance variance as before, and we contrasted trials in which participants had improved versus the ones in which they had not improved or got worse (see methods for details).

**Figure 3:**
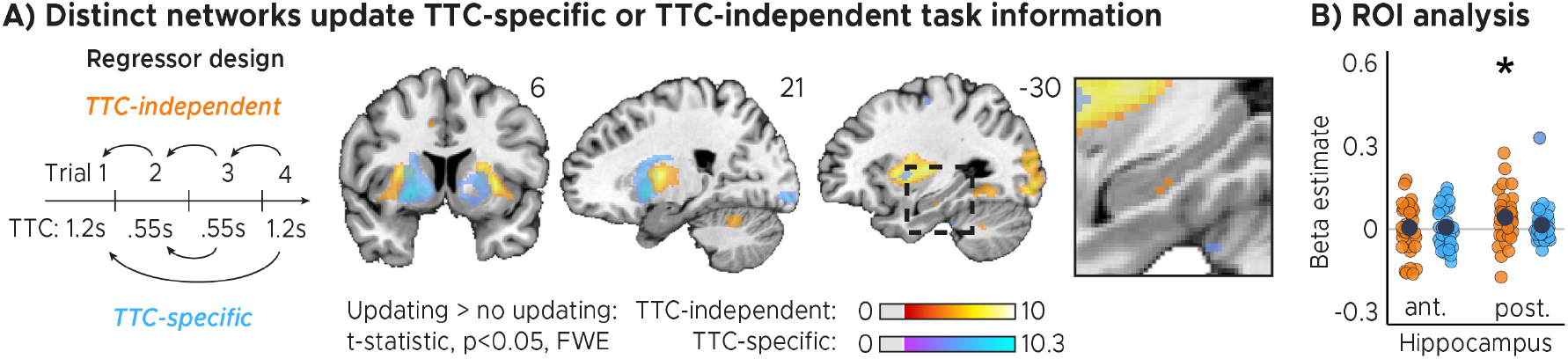
Distinct cortical and subcortical networks signal the updating of TTC-specific and TTC-independent task information. A) Left panel: Visual depiction of parametric modulator design. Two regressors per run modeled the improvement in behavioral performance since the last trial independent of the tested TTC (Regressor 1: TTC-independent) or the improvement since the last trial when the same target TTC was tested (Regressor 2: TTC-specific). Right panel: Voxel-wise analysis results for TTC-specific and TTC-independent regressors. We plot thresholded t-test results at 1mm resolution at p < 0.05 whole-brain Family-wise-error (FWE) corrected levels overlaid on a structural template brain. Insert zooming in on hippocampus and MNI coordinates added. B) Independent regions-of-interest analysis for the anterior (ant.) and posterior (post.) hippocampus. We plot the beta estimates obtained for TTC-independent in orange and TTC-specific regressors in blue. Depicted are the mean and SEM across participants (black dot and line) overlaid on single participant data as dots. Statistics reflect p<0.05 at Bonferroni-corrected levels (*) obtained using a group-level one-tailed one-sample t-test against zero.

We found both TTC-specific and TTC-independent activity throughout cortical and subcortical regions. Distinct areas engaged in either one or in both of these processes (Figs. 3A, S4). Crucially, we found that hippocampal activity signaled behavioral improvements independent of the TTC intervals tested. This effect was localized to the posterior section of the hippocampus (Fig. 3B, S2A; one-tailed one-sample *t* tests; TTC-independent: anterior HPC, *t*(33) = 0.36, *p* = 0.360, *p_fwe_* = 1, *d* = 0.06, *CI* : [−0.28,0.40], posterior HPC, *t*(33) = 2.81, *p* = 0.004, *p_fwe_* = 0.017, *d* = 0.48, *CI* : [0.12,0.85]; TTC-specific: anterior HPC, *t*(33) = 0.57, *p* = 0.285, *p_fwe_* = 1, *d* = 0.10, *CI* : [−0.24,0.44], posterior HPC, *t*(33) = 1.29, *p* = 0.103, *p_fwe_* = 0.413, *d* = 0.22, *CI* : [−0.12,0.57]). We then again estimated the functional connectivity profile of the hippocampal main effect using a PPI analysis (sphere with 4mm radius centered on the peak voxel at x=-30, y=-24, z=-18), revealing co-fluctuations in multiple regions including the putamen and the thalamus that were specific to behavioral improvements (Fig. S5).

These results suggest that the hippocampus updates information that is independent of the target TTC. This may support generalization performance by means of regularizing the encoded intervals based on the temporal context in which they were encoded. In our task, an efficient way of regularizing the encoded information is to bias one’s TTC estimates towards the mean of the TTC distribution, which corresponds to the regression effect that we observed on a behavioral level (Figs. 1B, S1C). Given the hippocampal feedback modulation and updating activity we reported above, we hypothesized that hippocampal activity should therefore also reflect the magnitude of the regression effect in behavior. To test this in a final analysis, we modeled the activity in each trial parametrically either as a function of performance (i.e. the absolute difference between estimated and true TTC) or as a function of the strength of the regression effect in each trial (i.e. the absolute difference between the estimated TTC and the mean of the tested intervals). Voxel-wise weights for these two regressors were estimated in two independent GLMs (see methods for details).

Our analyses showed that trial-wise hippocampal activity increased with better TTC-task performance (Figs. 4A, B; two-tailed one-sample *t* tests;anterior HPC, *t*(33) = −4.85, *p* = 2.9*x*10^-5^, *p_fwe_* = 5.8*x*10^-5^, *d* = −0.83, *CI* : [−1.24, −0.44]; posterior HPC, *t*(33) = −2.88, *p* = 0.007, *p_fwe_* = 0.014, *d* = −0.49, *CI* : [−0.86, −0.14]), and consistently also with the valence of the feedback received in the current trial (Fig. S3). In addition, however, and as predicted, it also reflected the trial-wise magnitude of the behavioral regression effect (Figs. 4A, B; two-tailed one-sample *t* tests; anterior HPC, *t*(33) = −5.55, *p* = 3.6*x*10^-6^, *p_fwe_* = 1.1*x*10^-5^, *d* = −0.95, *CI* : [−1.37, −0.55]; posterior HPC, *t*(33) = −1.06, *p* = 0.295, *p_fwe_* = 0.886, *d* = −0.18, *CI* : [−0.53, 0.16]). Activity in the anterior hippocampus was stronger in trials in which participants’ TTC estimates were more biased towards the mean of the sampled intervals (indicated by a negative beta estimate). Notably, similar effects were observed in prefrontal and posterior cingulate areas (Fig. 4A).

**Figure 4:**
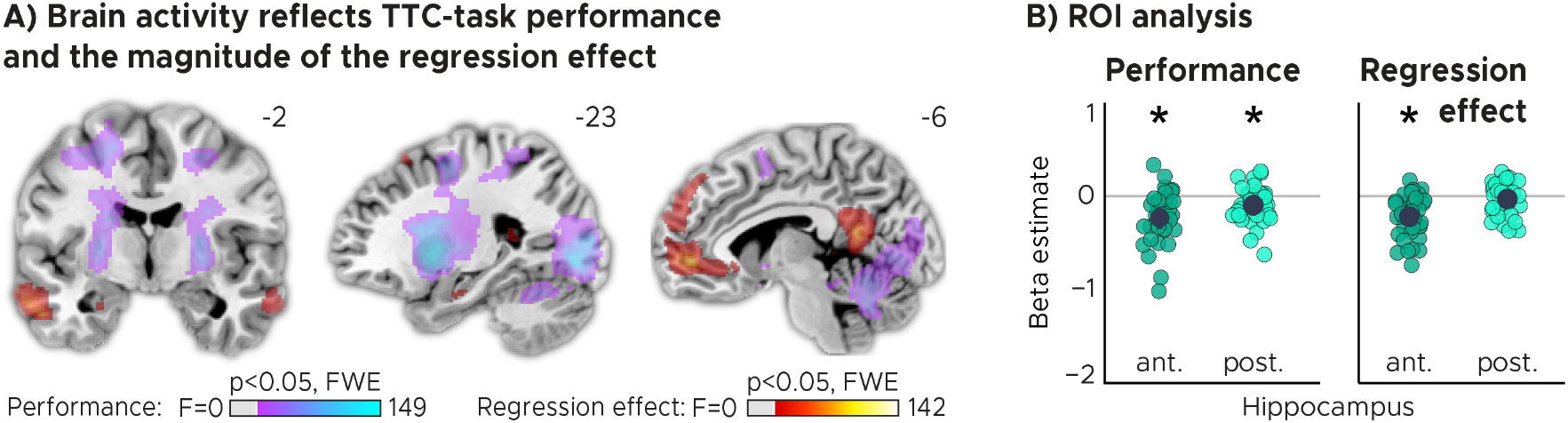
TTC-task performance vs. behavioral regression effect. A) Voxel-wise analysis. We plot thresholded F-test results for the task-performance regressor and the regression-to-the-mean regressor at 1 mm resolution overlaid on a structural template brain. MNI coordinates added. Distinct networks reflect task performance and the magnitude of the regression effect. B) Independent regions-of-interest analysis for the anterior (ant.) and posterior (post.) hippocampus. We plot the beta estimates obtained for each participant for each of the two regressors. Negative values indicate a linear increase between hippocampal activity and either task performance (left, Performance) or the magnitude of the regression effect (right, Regression effect). Depicted are the mean and SEM across participants (black dot and line) overlaid on single participant data (colored dots). Statistics reflect p<0.05 at Bonferroni-corrected levels (*) obtained using a group-level two-tailed one-sample t-test against zero.

### Eye tracking: no relevant biases in viewing behavior

To ensure that our results could not be attributed to systematic error patterns in viewing behavior, we analyzed the co-recorded eye tracking data of our participants in detail. After data cleaning (see methods), we used Kruskal-Wallis tests to control for differences in fixation accuracy across speed levels (Fig. S6A; *χ*^(2)^ = 0.61, *p* = 0.895, *ϵ*^2^ = 0.005, *CI* : [0.00,0.06]) and received-feedback levels (Fig. S6B; *χ*^(2)^ = 0.190, *p* = 0.909, *ϵ*^2^ = 0.002, *CI* : [0.00,0.10]). Moreover, we examined the relationship of the fixation error with TTC-task performance (Fig. S6C; Spearman’s *rho* = 0.17, *p* = 0.344) as well as with the behavioral regression effect (Fig. S6C; Spearman’s *rho* = 0.26, *p* = 0.131). None of these control analyses suggested that biased patterns in viewing behavior could hinder the interpretation of our results.

## Discussion

This study investigated how the brain extracts the statistical regularities of a sensorimotor timing task in a feedback-dependent manner. We specifically focused on the hippocampus, due to its known role in temporal coding and learning, asking how hippocampal processing may support behavioral flexibility, specificity and regularization. Moreover, we explored the larger brain-wide network involved in balancing these objectives. To do so, we monitored human brain activity with fMRI while participants estimated the time-to-contact between a moving target and a visual boundary. This allowed us to analyze brain activity as a function of task performance and as a function of the improvements in performance over time. We found that hippocampal activity as well as functional connectivity reflected the feedback participants received during this task, and its activity followed the performance improvements in a temporal-context-dependent manner. It signaled sensorimotor updating independent of the specific intervals tested, and its activity reflected trialwise behavioral biases towards the mean of the sampled intervals. In what follows, we discuss our results in the context of prior work on timing behavior and on hippocampal spatiotemporal coding. Moreover, we elaborate on the domain-general nature of hippocampal-cortical interactions and on the sensorimotor updating mechanisms that potentially underlie the effects observed in this study.

### Spatiotemporal coding in the hippocampus

The hippocampus encompasses neurons sensitive to elapsed time (Paton & Buonomano, 2018; Eichenbaum, 2014; Umbach et al., 2020). These cells might play an important role in guiding timing behavior (Nobre & van Ede, 2018), which potentially explains why hippocampal damage or inactivation impairs the ability to estimate durations in rodents (Meck et al., 1984) and humans (Richards, 1973). Our results are in line with these reports, showing that hippocampal fMRI activity also reflects participants’ TTC estimation ability (Fig. 4). They are also in line with other human neuroimaging studies suggesting that the hippocampus bridges temporal gaps between two stimuli during trace eyeblink conditioning (Cheng et al., 2008), and that it represents duration within event sequences (Barnett et al., 2014; Thavabalasingam et al., 2018, 2019). Our results speak to the above-mentioned reports by revealing that the hippocampus is an integral part of a widespread brain network contributing to sensorimotor updating of encoded intervals in humans (Figs. 2,3,4,S3,S4,S5). Moreover, they demonstrate a direct link between hippocampal activity, the feedback participants received and the behavioral improvements expressed over time (Fig. 3), emphasizing its role in feedback learning. Critically, the underlying process must occur in real-time when feedback is presented, suggesting that it plays out on short time scales. Notably, the human hippocampus is neither typically linked to sensorimotor timing tasks such as ours, nor is its activity considered to reflect temporal relationships on such short time scales. Instead, human hippocampal processing is often studied in the context of much longer time scales (Schiller et al., 2015; Eichenbaum, 2017), which showed that it may support the encoding of the progression of events into long-term episodic memories (Deuker et al., 2016; Montchal et al., 2019; Bellmund et al., 2021) or contribute to the establishment of chronological relations between events in memory (Gauthier et al., 2019, 2020). Intriguingly, the mechanisms at play may build on similar temporal coding principles as those discussed for motor timing (Yin & Troger, 2011; Eichenbaum, 2014; Howard, 2017; Palombo & Verfaellie, 2017; Nobre & van Ede, 2018; Paton & Buonomano, 2018; Bellmund et al., 2020, 2021; Shikano et al., 2021; Shimbo et al., 2021).

Ourtaskcan be solved by estimating temporal intervals directly, but also by extrapolating the movement of the fixation target over time, shifting the locus of attention towards the target boundary (Fig. 1). The brain may therefore likely monitor the temporal and spatial task regularities in parallel. Participants’ TTC estimates were further informed exclusively by the speed of the target, which inherently builds on tracking kinematic information over time, which may explain why TTC tasks also engage visual motion regions in humans (de Azevedo Neto & Amaro Júnior, 2018). While future studies could tease apart spatial and temporal factors explicitly, our results are in line with both accounts. For example, the hippocampus and surrounding structures represent maps of visual space in primates, which potentially mediate a coordinate system for planning behavior, integrating visual information with existing knowledge and to compute vectors in space (Nau et al., 2018; Bicanski & Burgess, 2020). These visuospatial representations are perfectly suited to guide attention and therefore the relevant behaviors in our task (Aly & Turk-Browne, 2017), which could be tested in the future akin to prior work using a similar paradigm (Nau et al., 2018a).

### The role of feedback in timed motor actions

Importantly, our results neither imply that the hippocampus acts as an “internal clock”, nor do we think of it as representing action sequences or coordinating motor commands directly. Rather, its activity may indicate the feedback-dependent updating of encoded information more generally and independent of the task that was used. The hippocampal formation has been proposed as a domain-general learning system (Kumaran, 2012; Schlichting & Preston, 2015; Chersi & Burgess, 2015; Schapiro et al., 2017; Wikenheiser et al., 2017; Behrens et al., 2018; Vikbladh et al., 2019; Geerts et al., 2020; Momennejad, 2020), which may encode the structure of a task abstracted away from our immediate experience. In contrast, the striatum was proposed to encode sensory states or actions, supporting the learning of task-specific (egocentric) information (Chersi & Burgess, 2015; Geerts et al., 2020). Together, the two regions may therefore play an important role in decision making in general also in other non-temporal domains.

Consistent with these ideas, we observed that striatal and hippocampal activity was modulated by behavioral feedback received in each trial (Figs. 2, S3). Similar feedback signals have been previously linked to learning (Schönberg et al., 2007; Cohen & Ranganath, 2007; Shohamy & Wagner, 2008; Foerde & Shohamy, 2011; Wimmer et al., 2012) and the successful formation of hippocampal-dependent long-term memories in humans (Wittmann et al., 2005). Moreover, hippocampal activity is known to signal learning in other tasks (Doeller et al., 2008; Foerde & Shohamy, 2011; Dickerson & Delgado, 2015; Wirth et al., 2009; Schapiro et al., 2017; Kragel et al., 2021). Here, we show a direct relationship between hippocampal activity and ongoing timing behavior, and we show that receiving behavioral feedback modulates widespread brain activity (Figs. 2, S3), which potentially reflects the involvement of these areas in the coordination of reward behavior observed earlier (LeGates et al., 2018). These regions include those serving sensorimotor functions, but also those encoding the structure of a task or the necessary value functions associated with specific actions (Lee et al., 2012).

The present study further demonstrates that activity in the hippocampus co-fluctuates with activity in other likely task-relevant regions in a task-dependent manner. We observed such co-fluctuations in the striatum and cerebellum, often associated with reward processing and action coordination (Bostan & Strick, 2018; Cox & Witten, 2019), the motor cortex, typically involved in action planning and execution, as well as the parahippocampal gyrus and medial parietal lobe, often associated with visual-scene analysis (Epstein & Baker, 2019). This may indicate that behavioral feedback also affects the functional connectivity profile of the hippocampus with those domain-selective regions that are currently engaged in the ongoing task. In the present report, this included the motor cortex, the parahippocampal gyrus, the medial parietal lobe and the cerebellum. This may also relate to previous reports of the cerebellum contributing temporal signals to cortical regions during similar tasks as ours (O’Reilly et al., 2008). Interestingly, we observed that hippocampal functional connectivity scaled negatively (Fig. 2C) with feedback valence, unlike its absolute activity, which scaled positively with feedback valence (Fig. 2A,B).

What might be the neural mechanism underlying sensorimotor updating signals in our task? Prior work has shown that hippocampal, frontal and striatal temporal receptive fields scale relative to the tested intervals, and that they re-scale dynamically when those tested intervals change (MacDonald et al., 2011; Gouvêa et al., 2015; Mello et al., 2015; Wang et al., 2018). This may enable the encoding and continuous maintenance of optimal task priors, which keep our actions well-adjusted to our current needs. We speculate that such receptive-field re-scaling also underlies the continuous updating effects discussed here, which likely build on both local and network-wide reweighting of functional connections between neurons and entire regions. Consistent with this idea and the present results, receptive-field re-scaling can occur on a trial-by-trial basis in the hippocampus (Shikano et al., 2021; Shimbo et al., 2021) but also in other regions such as the striatum (Mello et al., 2015; Gouvêa et al., 2015; Wang et al., 2018).

### A trade-off between specificity and regularization?

So far, we discussed how the brain may capture the temporal structure of a task and how the hippocampus supports this process. However, how do we encode specific task details while still forming representations that generalize well to new scenarios? In other words, how does the brain encode the probability distribution of the intervals we tested optimally without overfitting? Our behavioral and neuroimaging results suggest that this trade-off between specificity and regularization is governed by many regions, updating different types of task information in parallel (Fig. 3A). For example, hippocampal activity reflected performance improvements independent of the tested interval, whereas the caudate signaled improvements specifically over those trials in which the same TTC was tested. In the putamen, we found evidence for both processes (Fig. S4B). This suggests that different regions encode distinct task regularities in parallel to form optimal sensorimotor representations to balance specificity and regularization.

Notably, our results make a central prediction for future research. We anticipate that participants with stronger updating activity in the hippocampus should be able to generalize better to new scenarios, for example when new intervals are tested. While we could not test this prediction directly in our study, we did test for a link to a related phenomenon, and that is the regression effect we observed on the behavioral level. We found that TTC estimates regressed towards the mean of the sampled intervals in all participants (Figs. 1B, S1C), an effect that is well known in the timing literature (Jazayeri & Shadlen, 2010) and other domains (Petzschner & Glasauer, 2011; Petzschner et al., 2015). This regression effect likely reflects regularization in support of generalization (Roach et al., 2017), because time estimates are biased towards the mean of the tested intervals, and because the mean will likely be close to the mean of possible future intervals. We therefore hypothesized that this effect is grounded in the activity of the hippocampus, because it plays a central role in generalization in other non-temporal domains (Kumaran, 2012; Schlichting & Preston, 2015; Schapiro et al., 2017; Momennejad, 2020). Our analyses revealed that this was indeed the case. We found that hippocampal activity followed the magnitude of the regression effect in each trial (Fig. 4), potentially reflecting the temporal-context-dependent regularization of encoded intervals toward the grand mean of the tested intervals (Jazayeri & Shadlen, 2010).

In addition, our voxel-wise results showed that striatal subregions only tracked how accurate participants’ responses were, not how strongly they regressed towards the mean (Fig. 4A). This dovetails with literature on spatial-navigation (Doeller et al., 2008; Chersi & Burgess, 2015; Goodroe et al., 2018; Gahnstrom & Spiers, 2020; Geerts et al., 2020; Wiener et al., 2016), showing that the striatum supports the reinforcement-dependent encoding of locations relative to landmarks, whereas the hippocampus may help to encode the structure of the environment in a generalizable and map-like format. This matches the functional differences observed here in the time domain, where caudate activity reflects the encoding of individual details of our task such as the TTC intervals (Figs. 3A, S4A, B), while the hippocampus generalizes across TTCs to encode the overall task structure (Figs. 3A, B, S4A).

## Conclusion

In sum, we combined fMRI with time-to-contact estimations to show that the hippocampus sup-ports the formation of task-specific yet flexible and generalizable sensorimotor representations in real time. Hippocampal activity reflected trial-wise behavioral feedback and the behavioral improvements across trials, suggesting that it supports sensorimotor updating even on short time scales. The observed updating signals were independent from the tested intervals, and they explained the regression-to-the-mean biases observed on a behavioral level. This suggests that the hippocampus may encode temporal context in a behavior-dependent manner, and that it supports finding an optimal trade off between specificity and regularization. We show that it does so even in a fast-paced timing task typically considered to be hippocampal-independent. Our results show that the hippocampus supports rapid and feedback-dependent updating of sensorimotor representations, making it a central component of a brain-wide network balancing task specificity vs. regularization for flexible behavior in humans.

## Acknowledgements

We thank Raymundo Machado de Azevedo Neto for helpful comments on an earlier version of this manuscript. This work is funded by the European Research Council (ERC-CoG GEOCOG 724836 awarded to CFD). CFD’s research is further supported by the Max Planck Society, the Kavli Founda-tion, theJebsen foundation, the Centre of Excellence scheme of the Research Council of Norway - Centre for Neural Computation (223262/F50), The Egil and Pauline Braathen and Fred Kavli Centre for Cortical Microcircuits and the National Infrastructure scheme of the Research Council of Norway - NORBRAIN (197467/F50). RK’s research is supported by a CIDEGENT grant (CIDEGENT/2021/027) from the Valencian Community’s program for the support of talented researchers.

## Author Contributions

MN, IP and CFD developed the research questions. MN conceived the experimental idea. IP and MN designed the experimental paradigm, visualized the results and embedded them in the literature with help from RK, VW and CFD. IP implemented the experimental code and acquired and analyzed the data with close supervision and help from MN. MN wrote the manuscript with help from IP. CFD secured funding. RK, VW and CFD provided critical feedback and all authors discussed the results and edited the final manuscript. IP and MN are shared-first authors.

## Declaration of interest

The authors declare no conflicts of interest.

## Data and code availability

Source data and analysis code will be shared upon publication. Raw data are available from the authors upon request.

## Methods

### Participants

We recruited 39 participants for this study (16 females, 19-35 years old). Five participants were excluded: one participant did not comply with the task instructions; one was excluded due to a failure of the eye-tracker calibration; three participants were excluded due to technical issues during scanning. A total of 34 participants entered the analysis. The study was approved by the regional committee for medical and health research ethics (project number 2017/969) in Norway and participants gave written consent prior to scanning in accordance with the declaration of Helsinki (World Medical Association, 2013).

### Task

Participants performed two tasks simultaneously: a smooth pursuit visual-tracking task and a time-to-contact estimation task. The visual tracking task entailed fixation at a fixation disc that moved on predefined linear trajectories with one of four speeds: 4.2°/s, 5.8°/s, 7.5°/s and 9.1 °/s. Upon reaching the end of such a linear trajectory, the dot stopped moving until the second task was completed. This second task was a time-to-collision (TTC) estimation task in which participants indicated when the fixation target would have hit a circular boundary if it had continued moving. This boundary was a yellow circular line surrounding the target trajectory with 10° radius. Participants gave their response by pressing a button at the anticipated moment of collision. They performed this task while still keeping fixation, and the individual linear trajectories were all of the same length (10° visual angle), leading to four target TTC durations of 1.2s, 0.88s, 0.67s and 0.55s tested in a counter-balanced fashion across trials. After the button press, participants received feedback for 1 second informing them about the accuracy of their response. When participants *overestimated* the TTC, half of the fixation disc closest to the boundary changed color (orange or red) as a function of response accuracy (medium or low, respectively). When participants *underestimated* the TTC, half of the fixation disc further away from the boundary changed color. When participants were accurate, two opposing quadrants of the fixation disc would turn green. This allowed us to present feedback at fixation while keeping the number of informative pixels matched across feedback levels. To calibrate performance feedback across different TTC durations, the precise response window widths of each feedback level scaled with the speed of the fixation target. The following formula was used to scale the response window width: d ± ((k * d)/2) where d is the target TTC and k is a constant proportional to 0.3 and 0.15 for high and medium accuracy, respectively. This ensured that participants received approximately the same feedback for tested TTCs despite the known differences in absolute performance between target TTCs due to inherent scalar variability (Gibbon, 1977). When no response was given, participants received low-accuracy feedback (two opposing quadrants of the fixation dot turned red) after a 4 seconds timeout. After the feedback, the disc remained in its last position for a variable inter-trial interval (ITI) sampled randomly from a uniform distribution between 0.5 seconds and 1.5 seconds. Following the end of the ITI, the dot continued moving in a different direction. In the course of 768 trials, each target TTC was sampled 192 times. We sampled eye-movement directions with 15° resolution, leading to an overall trajectory that was star-shaped, similar to earlier reports (Nau et al., 2018a). The full trajectory was never explicitly shown to the participants.

### Behavioral analysis

Participants indicated the estimated TTC in each trial via button press. In line with previous work (Jazayeri & Shadlen, 2010), participants tended to overestimate shorter durations and underestimate longer durations (Fig. 1 B). In order to quantify this behavioral effect we extracted the slope value of a linear regression line fit between estimated and target TTCs separately for each participant. A slope of 1 would indicate that participants performed perfectly accurately for all intervals. A slope of 0 would indicate that participants always gave the same response independent of the tested interval, fully regressing to the mean of the sampled intervals. Two separate one-tailed one-sample *t* tests (against 1 or 0) were performed to corroborate that participants’ slope values regressed towards the mean of the sampled TTCs (Fig. S1C). A Spearman’s rank-order correlation tested if slope values correlated with the percent of high accuracy trials (Fig. S1D), to further demonstrate that participants relied to different degrees on both, the target TTCs and the mean of the sampled TTCs, in order to achieve an optimal performance tradeoff. As the TTC task progressed, it would be expected that participants adjusted their TTC estimates in order to find the best tradeoff. Thus, we tested if the slope converged over time towards the value of 0.5 (the slope value between veridical performance and the mean of the sampled TTCs) by using a linear mixed-effects model with task segment as a predictor, the absolute difference between the slope and the value of 0.5 as the dependent variable and participants as the error term (Fig. S1E). As a measure of behavioral performance, we computed the absolute TTC-error defined as the absolute difference in estimated and true TTC for each target-TTC level. Participants received feedback after each trial corresponding to the absolute TTC error of that trial. On average, 46.9% (*σ* = 9.1) of trials were of *high accuracy*, 31.2% (*σ* = 3.9) were of *medium* accuracy and 21.1% (*σ* = 9.8) were of *low* accuracy (Fig. 1C). Moreover, we found that this feedback distribution was indeed similar across target-TTC levels as planned (Fig. S1B). To control that there was no systematic and predictable relationship between subsequent trials on a behavioral level, we estimated the n-1 Pearson autocorrelation between feedback values received on each trial and then performed a two-tailed one-sample t-test on group level against zero using the extracted correlation coefficients from each participant (Fig. S1A). To further test participants’ performance improvements over time, we used a linear mixed-effects model with task segment as predictor, absolute TTC-error as the dependent variable and participants as the error term (Fig. S1 F).

### Imaging data acquisition & preprocessing

Imaging data were acquired on a Siemens 3T MAGNETOM Skyra located at the St. Olavs Hospital in Trondheim, Norway. A T1-weighted structural scan was acquired with 1mm isotropic voxel size. Following EPI-parameters were used: voxel size=2mm isotropic, TR=1020ms, TE=34.6ms, flip angle=55°, multiband factor=6. Participants performed a total of four scanning runs of 16-18 minutes each including a short break in the middle of each run. Functional images were corrected for head motion and co-registered to each individual’s structural scan using SPM12 (www.fil.ion.ucl.ac.uk/spm/). We used the FSL topup function to correct field distortions based on one image acquired with inverted phase-encoding direction (https://fsl.fmrib.ox.ac.uk/fsl/fslwiki/topup). Functional images were then spatially normalized to the Montreal Neurological Institute (MNI) brain template and smoothed with a Gaussian kernel with full-width-at-half-maximum of 4 mm for regions-of-interest analysis or with 8 mm for whole-brain analysis. Time series were high-pass filtered with a 128 s cut-off period. The results of all voxel-wise analyses were overlaid on a structural T1-template (colin27) of SPM12 for visualization.

### Regions of interest definition and analysis

Regions-of-interest masks for different brain areas were generated for each individual participant based on the automatic parcellation derived from FreeSurfer’s structural reconstruction (https://surfer.nmr.mgh.harvard.edu/). The ROIs used in the present study include the Hippocampus as main area of interest (Fig. S2A) as well as the Caudate Nucleus, Nucleus Accumbens, Thalamus, Putamen, Amygdala and Globus Pallidum (Fig. S2B). The hippocampal ROI was manually segmented following previous reports into its anterior and posterior sections based on the location of the uncal apex in the coronal plane as a bisection point (Poppenk et al., 2013). We did this because prior work suggested functional differences between anterior and posterior hippocampus with respect to their contributions to memory-guided behavior (Poppenk et al., 2013). All individual ROIs were then spatially normalized to the MNI brain template space and re-sliced to the functional imaging resolution using SPM12. All ROI analyses were conducted using 4mm spatial smoothing.

All ROI analyses described in the following were conducted using the following procedure. We extracted beta estimates estimated for the respective regressors of interest for all voxels within a region in both hemispheres, averaged them across voxels within that region and hemispheres and performed one-sample t-tests on group level against zero as implemented in the software R (https://www.R-project.org).

### Brain activity as a function of performance feedback on the previous trial

To examine how feedback modulates activity in the subsequent trial, we used a mass-univariate general linear model (GLM) analysis to model the activity of each voxel and trial as a function of feedback received in the previous trial. The GLM included three regressors modeling the feedback levels, one for ITIs, one for button presses and one for periods of rest, which were all convolved with the canonical hemodynamic response function of SPM12. In addition, the model included the six realignment parameters obtained during pre-processing as well as a constant term modeling the mean of the time series. On the group level, we then contrasted the weights obtained for the low error vs. high error regressors and tested for differences using t-tests implemented in SPM12 (Fig. 2A).

Additionally, we again conducted ROI analyses for the anterior and posterior sections of the hippocampus (Fig. S2A) following the same procedure as described earlier (section “Regions of interest definition and analysis”). Here, we tested beta estimates obtained in the first-level analysis for the feedback-in-previous-trial regressor of interest (Fig. 2B).

### Hippocampal functional connectivity as a function of previous-trial performance feedback

We conducted a psychophysiological interactions (PPI) analysis to examine whether hippocampal functional connectivity with the rest of the brain depended on the participant’s performance on the previous trial. To do so, we centered a sphere onto the group-level peak effects within the HPC using main-effect GLM described in the previous section. The sphere was 4mm in radius and was centered on the following MNI coordinates: x=-32, y=-14, z=-14. The GLM included a PPI regressor, a nuisance regressor accounting for the main effect of past-trial performance, and a nuisance regressor explaining variance due to inherent physiological signal correlations between the HPC and the rest of the brain. The PPI regressor was an interaction term containing the element-by-element product of the task time course (effects due to past-trial performance) and the HPC spherical seed ROI time course. The estimated beta weight corresponding to the interaction term was then tested against zero on the group-level using a t-test implemented in SPM12 (Fig. 2C). This revealed brain areas whose activity was co-varying with the hippocampus seed ROI as a function of past-trial performance (n-1).

### Brain activity as a function of current-trial performance feedback

We used a GLM to analyze the time courses of all voxels in the brain as a function of feedback received at the end of each trial. The model included one mean-centered parametric modulator per run with three levels reflecting the feedback received in each trial. The feedback itself was a function of TTC error in each trial (high accuracy = 0, medium accuracy = 0.5 and low accuracy = 1). In addition, we added three nuissance regressors per run modeling ITIs, button presses, and periods of rest. These regressors were convolved with the canonical hemodynamic response function of SPM12. Moreover, the realignment parameters and a constant term were again added. We estimated weights for all regressors and conducted a t-test against zero using SPM12 for our feedback regressors of interest on the group level (Fig. S3A). Importantly, positive t-scores indicate a positive relationship between fMRI activity and TTC error and hence with poor behavioral performance. Conversely, negative t-scores indicate a negative relation between the two variables and hence better behavioral performance.

In addition to the voxel-wise whole-brain analyses described above, we conducted independent ROI analyses for the anterior and posterior sections of the hippocampus (Fig. S2A). Here, we tested the beta estimates obtained in our first-level analysis for the feedback regressor of interest (Fig. S3B). See section “Regions of interest definition and analysis” for more details.

### Brain activity as a function of improvements in behavioral performance across trials

We used a GLM to analyze activity changes associated with behavioral improvements across trials. One regressor modelled the main effect of the trial and two parametric regressors modeled the following contrasts: trials in which behavioral performance improved *vs*. trials in which behavioral performance did not improve or got worse relative to the previous trial. These regressors modeled the behavioral improvements either relative to the previous trial, and therefore independently of TTC (likely serving regularization), or relative to the previous trial in which the same target TTC was presented (likely serving specificity). These two regressors reflect the tests for target-TTC-independent and target-TTC-specific updating, respectively. Improvement in performance was defined as receiving feedback of higher valence than in the corresponding previous trial. The same nuisance regressors were added as in the other GLMs and all regressors except the realignment parameters and the constant term were convolved with the canonical hemodynamic response function of SPM12. On the group level, we tested the two parametric regressors of interest against zero using a t-test implemented in SPM12, effectively contrasting trials in which behavioral performance improved against trials in which behavioral performance did not improve or got worse relative to the respective previous trials (Fig. 3A). All runs were modeled separately.

Moreover, we again conducted ROI analyses for the anterior and posterior sections of the hippocampus (Fig. S2A) following the same procedure as described earlier (see section “Regions of interest definition and analysis”). Here, we tested beta estimates obtained in the first-level analysis for the TTC-specific and TTC-independent updating regressors using one-tailed one-sample t-tests (Fig. 3B). In addition, to test which specific subcortical regions were involved in these processes, we conducted post-hoc ROI analyses for subcortical regions after the whole-brain results were known (Fig. S4B; one-tailed one-sample *t* tests; TTC-specific: caudate: *t*(33) = 5.95, *p* = 5.6*x*10^-7^, *p_fwe_* = 3.4*x*10^-6^, *d* = 1.02, *CI* : [0.61,1.45], nucleus accumbens: *t*(33) = 4.41, *p* = 5.2*x*10^-5^, *p_fwe_* = 3.1*x*10^-4^, *d* = 0.76, *CI* : [0.38,1.15], globus pallidus: *t*(33) = 7.05, 2.3*x*10^-8^, *p_fwe_* = 1.4*x*10^-7^, *d* = 1.21, *CI* : [0.77,1.67], putamen: *t*(33) = 8.07, *p* = 1.3*x*10^-9^, *p_fwe_* = 7.7*x*10^-9^, *d* = 1.38, *CI* : [0.92,1.88], amygdala: *t*(33) = 1.78, *p* = 0.042, *p_fwe_* = 0.255, *d* = 0.30, *CI* : [−0.04,0.66], thalamus: *t*(33) = 2.61, *p* = 0.007, *p_fwe_* = 0.007, *d* = 0.45, *CI* : [0.09,0.81]; TTC-independent: caudate: *t*(33) = −0.67, *p* = 0.746, *p_fwe_* = 1, *d* = −0.11, *CI* : [−0.46,0.23], nucleus accumbens: *t*(33) = 1.82, *p* = 0.039, *p_fwe_* = 0.235, *d* = 0.31, *CI* : [−0.04,0.66], globus pallidus: *t*(33) = 7.06, *p* = 2.2*x*10^-8^, *p_fwe_* = 1.3*x*10^-7^, *d* = 1.21, *CI* : [0.77,1.68], putamen: *t*(33) = 6.21, *p* = 2.6*x*10^-7^, *p_fwe_* = 1.6*x*10^-6^, *d* = 1.06, *CI* : [0.65,1.50], amygdala: *t*(33) = 4.25, *p* = 8.3*x*10^-5^, *p_fwe_* = 4.9*x*10^-4^, *d* = 0.73, *CI* : [0.35,1.12], thalamus: *t*(33) = 4.05, *p* = 1.5*x*10^-4^, *p_fwe_* = 8.9*x*10^-4^, *d* = 0.69, *CI* : [0.32, 1.08]). The subcortical ROIs (Fig. S2B) were based on the FreeSurfer parcellation as described in the section “Regions of interest definition and analysis”.

### Hippocampal functional connectivity as a function of TTC-independent updating

To examine which brain regions whose activity co-fluctuated with the one of the hippocampus during TTC-independent updating, we again conducted a PPI analysis similar to the one described earlier. A spherical seed ROI with a radius of 4 mm was centered around the hippocampal group-level peak effect (x=-30, y=-24, z=-18) observed for the TTC-independent updating regressor described above. The GLM included a PPI regressor and two nuisance regressors accounting for task-related effects from our contrast of interest (Behavioral improvements vs. no behavioral improvements) as well as physiological correlations that could arise due to anatomical connections to the hippocampal seed region or shared subcortical input. On the group-level, we then tested the weights estimated for our PPI regressor of interest against zero using a t-test implemented in SPM12. This revealed areas whose activity co-fluctuated with the one of the hippocampus as a function TTC-independent updating (Fig. S5A).

Moreover, we conducted independent ROI analyses for subcortical regions as described in the section “Regions of interest definition and analysis”. Here, we tested the beta estimates obtained for the hippocampal seed-based PPI regressor of interest (Fig. S5B; one-tailed one-sample *t* tests: caudate: *t*(33) = 1.06, *p* = 0.149, *p_fwe_* = 0.894, *d* = 0.18, *CI* : [−0.16,0.53], putamen: *t*(33) = 2.79, *p* = 0.004, *p_fwe_* = 0.026, *d* = 0.48, *CI* : [0.12,0.84], globus pallidus: *t*(33) = 2.52, *p* = 0.008, *p_fwe_* = 0.050, *d* = 0.43, *CI* : [0.08,0.79], amygdala: *t*(33) = 2.60, *p* = 0.007, *p_fwe_* = 0.041, *d* = 0.45, *CI* : [0.09,0.81], nucleus accumbens: *t*(33) = −1.14, *p* = 0.869, *p_fwe_* = 1, *d* = −0.20, *CI* : [−0.54,0.15], thalamus: *t*(33) = 2.71, *p* = 0.005, *p_fwe_* = 0.032, d = 0.46, *CI* : [0.11,0.83]).

### Brain activity as a function of behavioral performance and as a function of the behavioral regression effect

To examine the neural underpinnings governing specificity and regularization in timing behavior in detail, we analyzed the trial-wise activity of each voxel as a function of performance in the TTC task (i.e. the absolute difference between estimated and true TTC in each trial) and as a function of the regression effect in behavior (i.e. the absolute difference between the estimated TTC and the mean of the sampled intervals, which was 0.82 s). To avoid effects of potential co-linearity between these regressors, we estimated model weights using two independent GLMs, which modeled the time course of each trial with either one of the two regressors. In addition, we again accounted for nuisance variance as described before, and all regressors except the realignment parameters and the constant term were convolved with the canonical HRF of SPM12. After fitting the model, we used the weights estimated for the two regressors to perform voxel-wise F-tests using SPM12, revealing activity that was correlated with these two regressors independent of the sign of the correlation (Fig. 4A). In addition, we again performed ROI analyses using two-tailed one-sample t-tests for the anterior and posterior hippocampus (Figs. S2A, 4B).

### Eye tracking: Fixation quality does not affect the interpretation of our results

We used an MR-compatible infrared eye tracker with long-range optics (Eyelink 1000) to monitor gaze position at a rate of 500 hz during the experiment. After blink removal, the eye tracking data was linearly detrended, median centered, downsampled to the screen refresh rate of 120 hz and smoothed with a running-average kernel of 100 ms. Kruskal-Wallis tests were used in order to test for potential biases in fixation error across speeds (Fig. S6A) or across feedback levels (Fig. S6B). Moreover, we tested if differences in fixation error could either explain individual differences in the regression effect, or individual differences in absolute TTC error in behavior using Spearman’s rank-order correlations (Fig. S6C).

## Supplementary Material

**Figure S1:**
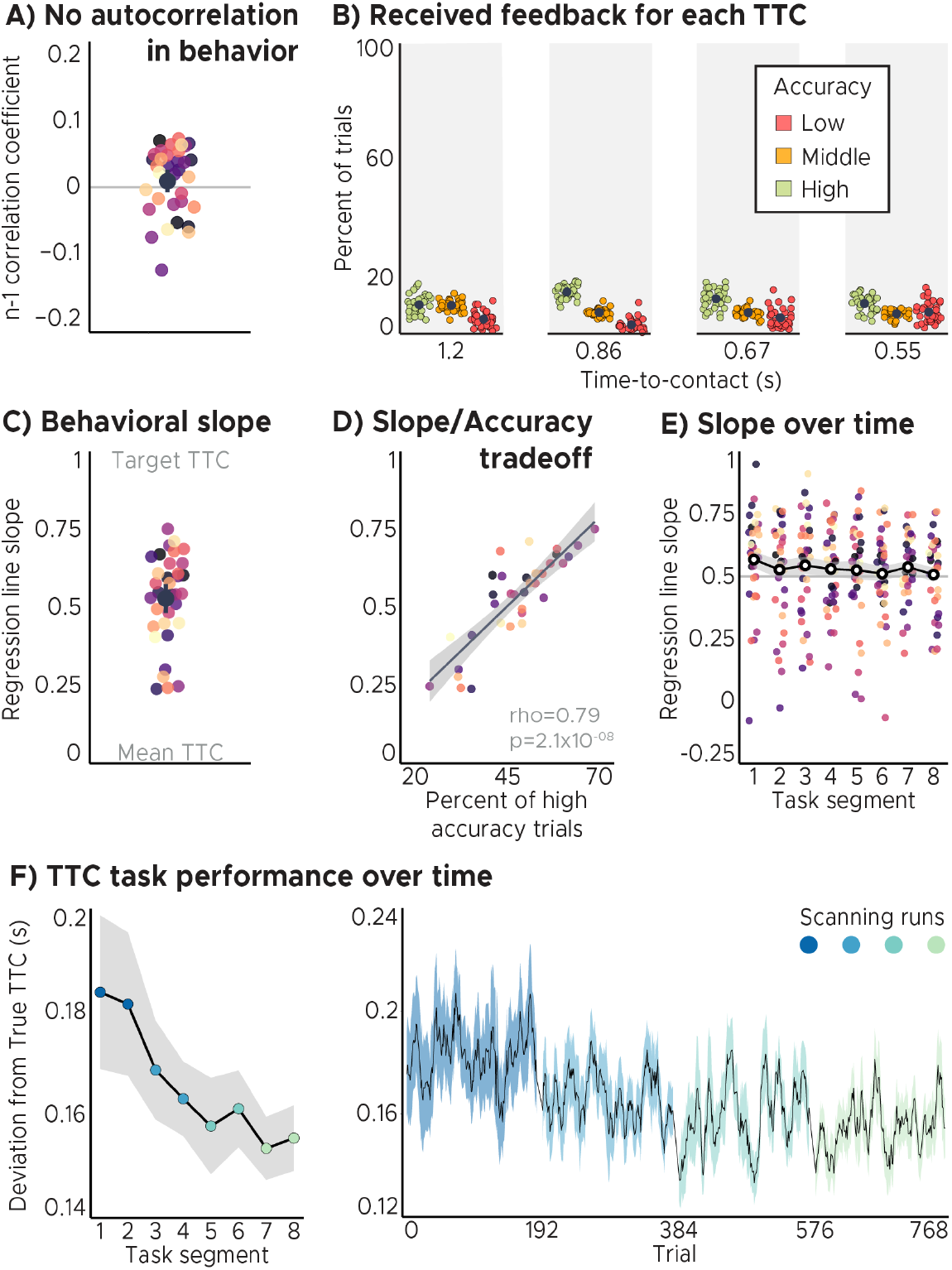
Behavioral analyses. A) No autocorrelation in the behavioral feedback over trials. The feedback in one trial did not predict feedback in the following trial. Displayed values correspond to the Pearson n-1 correlation coefficient. B) Feedback distributions for all speed levels. Participant’s received approximately the same feedback for all speed levels/target TTCs. C) Behavioral regression effect. We plot linear regression-line slopes predicting estimated TTCs as a function of target TTCs for each participant. A slope of 1 indicates perfect performance. A slope of 0 indicates that participants always gave the same response independent of the target TTC. We found that the slope coefficients clustered at around 0.5, suggesting that participants’ responses were biased towards the mean of the sampled intervals. ABC) Depicted are the mean and SEM across participants (black dot and line) overlaid on single participant data (dots). D) Performance trade-off between the regression effect and TTC accuracy. Participants with higher TTC accuracy exhibited a weaker regression effect, reflected in larger regression-line slopes (same data as in C). Each dot represents a single participant. Regression line (black) and SEM (gray shade) were added. E) Behavioral regression effect over time. Participants’ slope coefficients converged towards the value of 0.5 as they progressed through the task. Each dot represents a single participant. Mean (black and white dot) and SEM (gray shade) were added. ACDE) Participants were color coded. F) TTC task performance over time. Left panel: Across-trial-average performance over task segments. Right panel: task performance over trials. We plot the mean (black line) and SEM (shaded area) across participants. Run identity color coded. Participants’ task performance improved over time.

**Figure S2:**
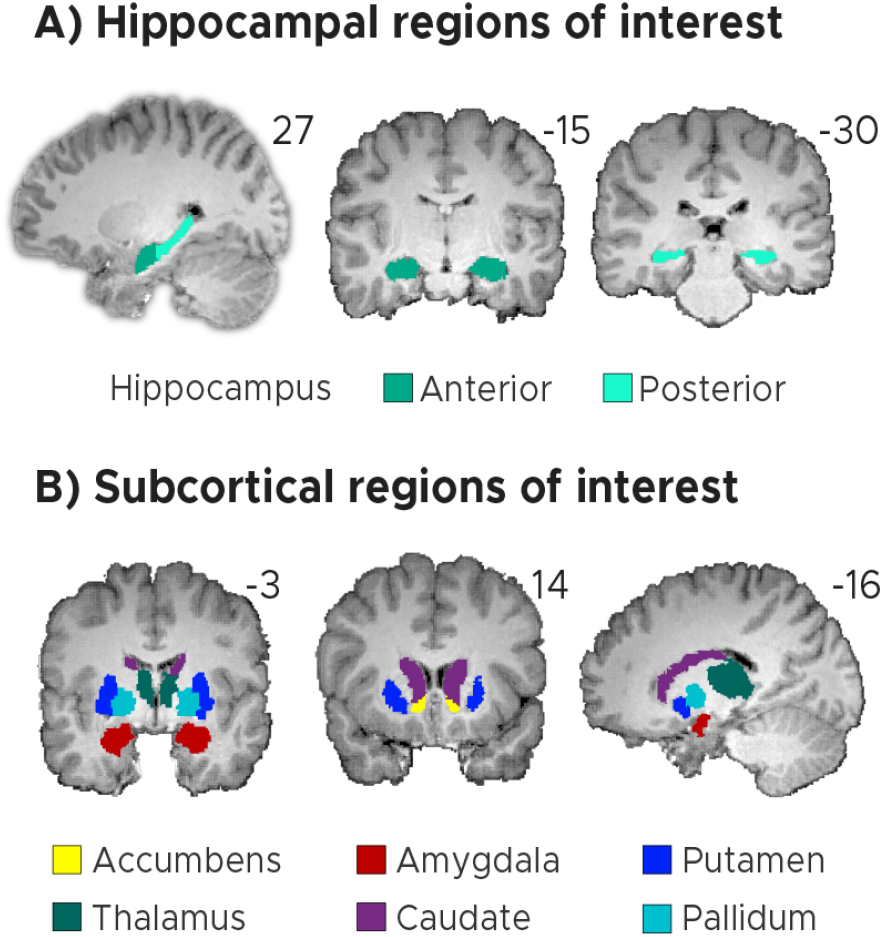
Regions of interest (ROIs). A) Anterior and posterior hippocampal ROIs. B) Subcortical regions-of-interest (ROIs) for the nucleus accumbens, the amygdala, the thalamus, the caudate, the putamen and the pallidum. AB) ROIs shown for a sample participant superimposed onto the skull-stripped structural T1-scan of that participant. These masks were created using FreeSurfer’s cortical and subcortical parcellation.

**Figure S3:**
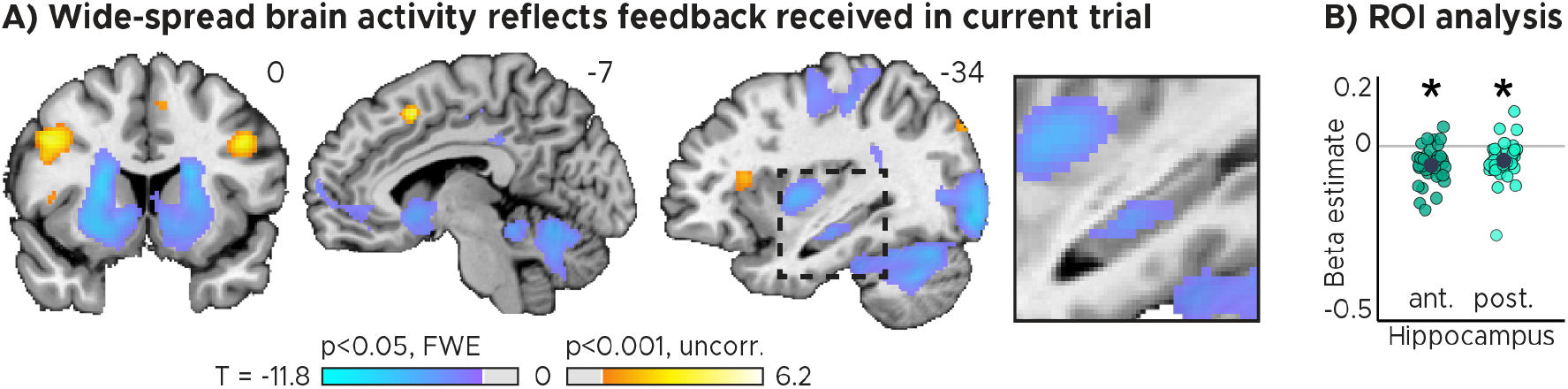
Brain regions signalling behavioral feedback in current trial. Activity in each trial was modeled parametrically as a function of the feedback received at the end of the trial. A) Voxel-wise analysis. We plot thresholded t-test results at 1 mm resolution overlaid on a structural template brain. MNI coordinates and insert zooming in on the hippocampus added. A large network of regions signalling TTC performance included the hippocampus, striatum and cerebellum. B) Independent regions-of-interest analysis for the anterior (ant.) and posterior (post.) hippocampus. We plot the beta estimate obtained for the parametric modulator modeling trial-wise activity as a function of task performance. Negative values indicate that smaller errors, and higher-accuracy feedback, led to stronger activity. Depicted are the means and SEM across participants (black dot and line) overlaid on single participant data (coloured dots). Statistics reflect p < 0.05 at Bonferroni-corrected levels (*) obtained using a group-level two-tailed one-sample t-test against zero.

**Figure S4:**
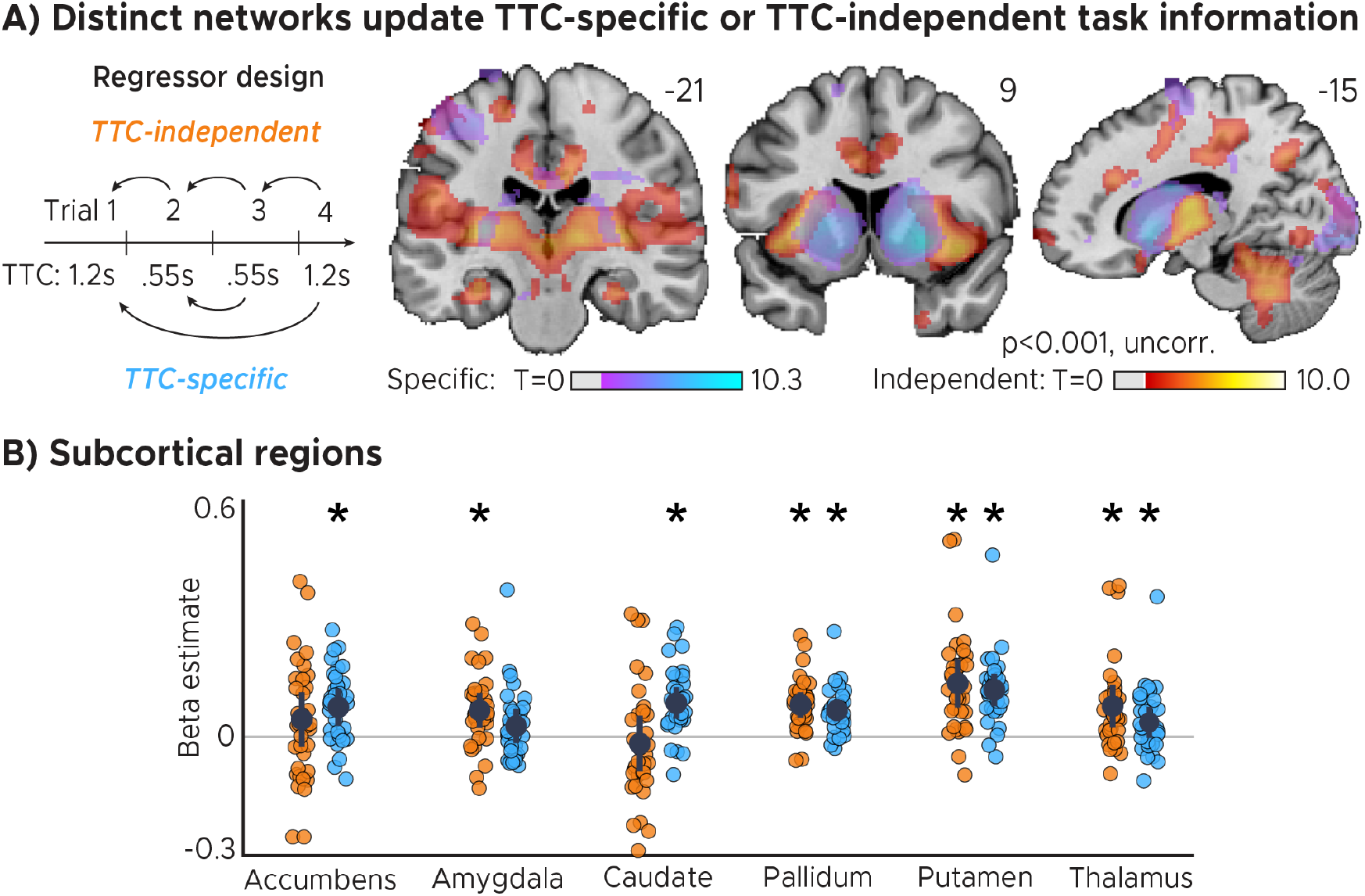
Distinct networks support TTC-specific and TTC-independent updating. A) Voxel-wise mass-univariate GLM results for TTC-independent and TTC-specific parametric regressors. We plot thresholded t-test results at 1mm resolution. Activity maps were overlaid on a structural template brain. Positive t-scores indicate a relationship between brain activity and the updating of either TTC-specific or TTC-independent information respectively. B) ROI-analysis results for subcortical regions for TTC-independent (orange dots) and TTC-specific regressors (blue dots). Depicted are the mean and SEM across participants (black dot and line) overlaid on single participant data. Statistics reflect p<0.05 at Bonferroni-corrected levels (*) obtained using a group-level one-tailed one-sample t-test against zero.

**Figure S5:**
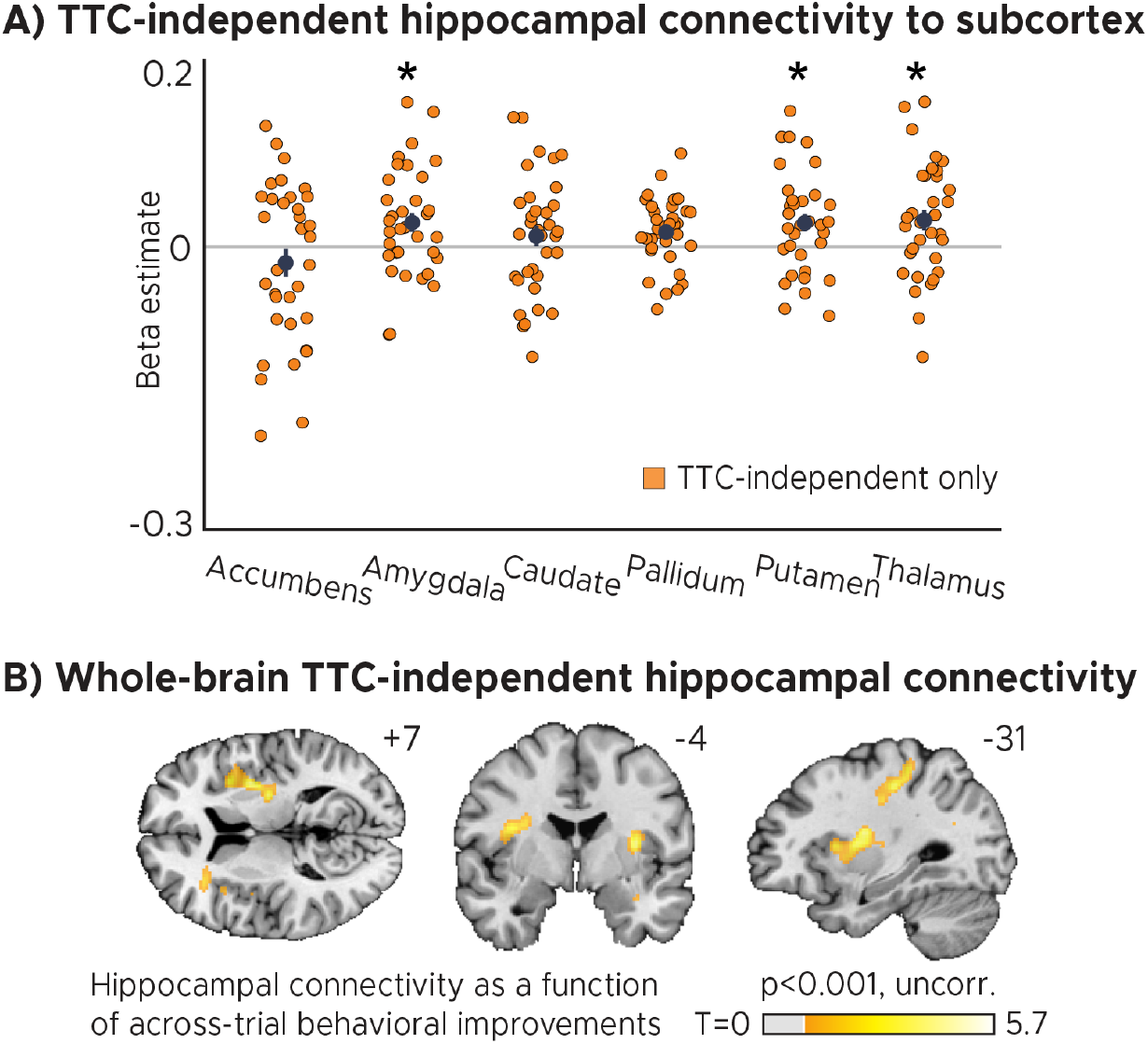
TTC-independent hippocampal connectivity. A) Regions of interest analysis for subcortical regions estimated using a Psychophysiological-interactions (PPI) analysis conducted using the hippocampal effect in Fig. 3A as a seed. Positive beta estimates indicate that functional connectivity between each ROI and the hippocampal seed depended on how much participants TTC-task performance improved across trials. Depicted are the mean and SEM across participants (black dot and line) overlaid on single participant data for the nucleus accumbens, the amygdala, the caudate, the globus pallidum, the putamen and the thalamus. Statistics reflect p<0.05 at Bonferroni-corrected levels (*) obtained using a group-level one-tailed one-sample t-test against zero. B) Whole-brain voxel-wise t-test results for the TTC-independent hippocampal connectivity overlaid on a structural template brain at 1mm resolution. MNI coordinates added.

**Figure S6:**
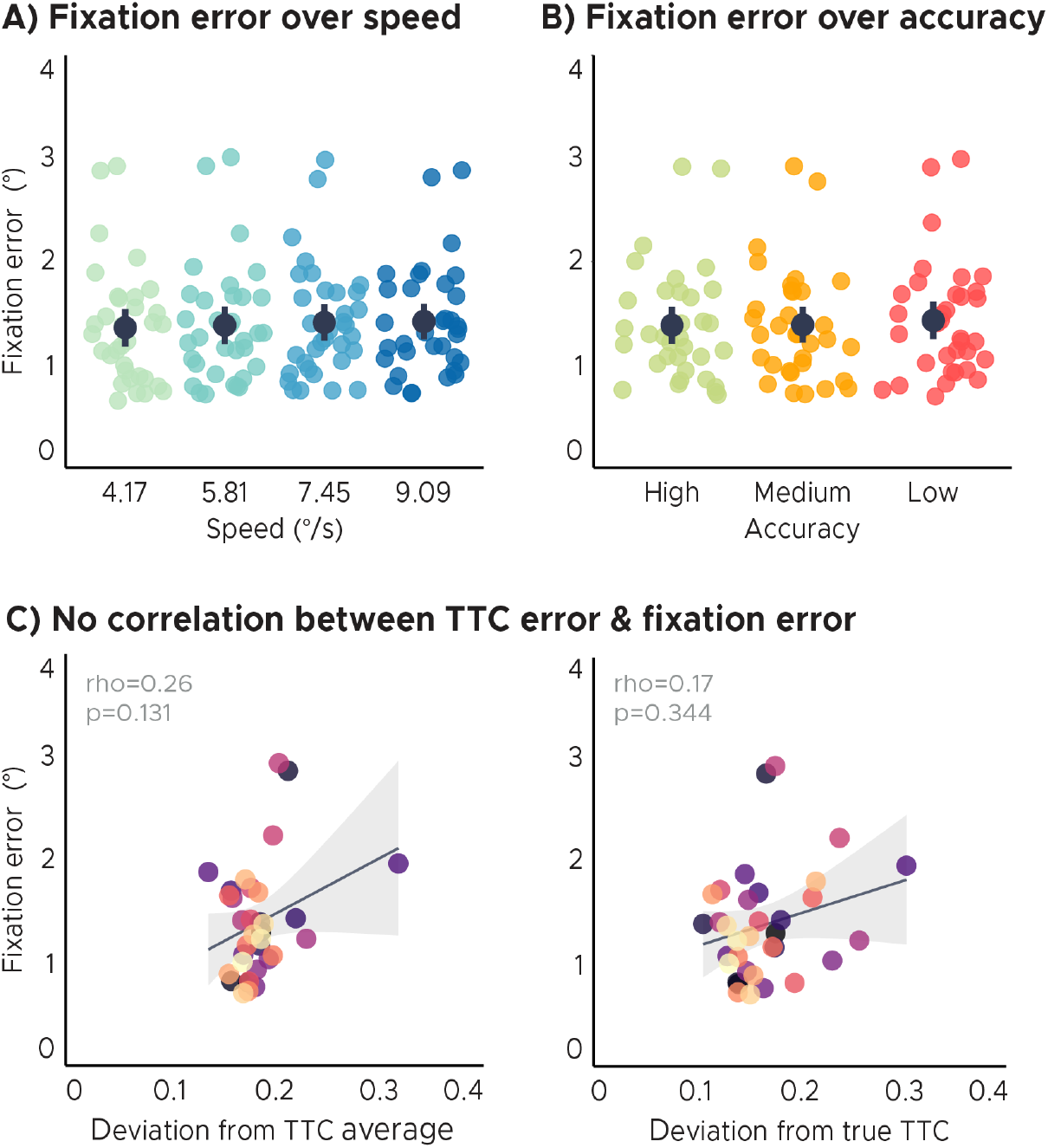
Eye tracking analyses. A) Fixation error over speed. There were no significant differences in fixation error across speed levels/target TTC’s. B) Fixation error over TTC-task accuracy. There were no significant differences in fixation error across TTC-task accuracy levels. C) No correlation of the behavioral regression-to-the-mean effect or TTC-task performance with fixation error. Fixation quality does not affect the interpretation of the imaging results presented in this study. Each dot represents a single participant. Participants were color coded. Regression line (black) and standard error (gray shade). AB) Group-level mean and SEM depicted as black dot and line overlaid on single participant data.

## Notes

### Competing Interest Statement

The authors have declared no competing interest.

